# Loop extrusion driven volume phase transition of entangled chromosomes

**DOI:** 10.1101/2022.03.18.484867

**Authors:** Tetsuya Yamamoto, Helmut Schiessel

## Abstract

Mitotic chromosomes without nucleosomes have been reconstituted in recent experiments. When topo II is depleted from the reconstituted chromosomes, these chromosomes are entangled and form ‘sparklers’, where DNA is condensed in the core with linker histone H1.8 and condensin is localized at the periphery. To understand the mechanism of the assembly of sparklers, we here take into account the loop extrusion by condensin in an extension of the theory of entangled polymer gels. The loop extrusion stiffens an entangled DNA network because DNA segments in the elastically effective chains are translocated to loops, which are elastically ineffective. Our theory predicts that the loop extrusion by condensin drives the volume phase transition that collapses a swollen entangled DNA gel as the stiffening of the network destabilizes the swollen phase. This is an important element to understand the mechanism of the assembly of the reconstituted chromosomes.

## Introduction

At the entry of mitosis, chromatin, which has been distributed within the nucleus in interphase, is condensed into mitotic chromosomes by metaphase via a series of structural transitions. A mitotic chromosome is composed of two sister chromatids, which are bound at the centromere by cohesin. Each sister chromatid forms a cylindrical structure, where the loops of chromatin are densely packed along the axis. This structure has attracted molecular and cell biologists for more than 160 years.^1^ A crucial player in the formation of mitotic chromosomes is condensin, a ring-shaped protein complex that is loaded onto DNA by embracing it.^2^ The loop extrusion theory predicts that the chromatin loops of mitotic chromosomes are produced by condensin via the loop extrusion process, with which chromatin segments are uni-directionally transported from the chromosome axis to the loops.^3–6^ The loop extrusion activity of condensin was visualized and characterized by using the DNA curtain technique.^7–9^ Theoretically, mitotic chromosomes are studied by using the analogy with bottle brush polymers, in which side chains are densely packed along their main chains.^10,11^ In recent experiments Hirano and coworkers reconstituted mitotic chromosomes.^12–14^ They showed that condensin and topoisomerase II (topo II) are necessary components to assemble mitotic chromosomes for DNA derived from mouse sperms by depleting protamines and nucleosomes, ^13^ see fig. 1**a**. Topo II is an enzyme that disentangles DNA by transiently breaking the double-strand and then ligating it. When both topo II and condensin are depleted, chromosomes behave as swollen entangled networks, failing to be individualized and to form the condensed cylindrical structure, see fig. 1**b**. Shintomi and Hirano have recently discovered that when condensin is present and topo II is depleted, DNA forms a new structure, called sparkler, ^14^ see fig. 1**c**. The feature of the sparkler is that DNA and histone H1.8 are condensed in the core and condensin I is excluded to the periphery.

**Figure 1:**
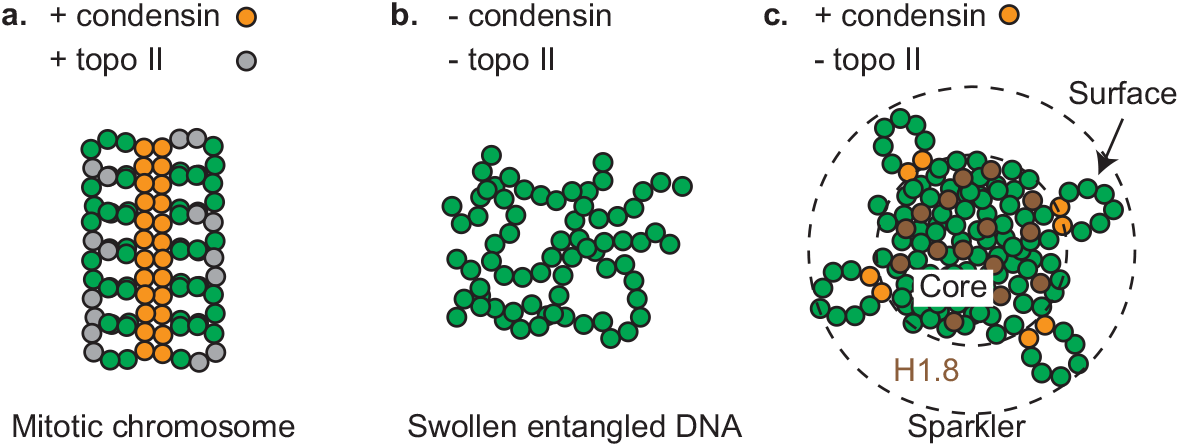
Polymorphism of reconstituted chromosomes, from which nucleosomes are removed. **a**. With condensin and topo II, the reconstituted chromosomes form a structure akin of a mitotic chromosome. **b**. Without condensin and topoisomerase II (topo II), the reconstituted chromosomes behave as an entangled gel. **c**. When only topo II is depleted, the reconstituted chromosomes form sparklers, where DNA and linker histone H1.8 are condensed in the core and condensin is localized at the periphery. DNA segments, condensin, topo II, and H1.8 are represented by green, orange, gray, and brown circles.

To understand the mechanism of the assembly of sparklers, it is instructive to think of how the loop extrusion by condensin affects the mechanics of entangled DNA networks. In polymer physics, entanglements are treated as effective crosslinks that can slide along the chains (so-called slip-links). ^15–18^ Networks of synthetic polymers are prepared by crosslinking polymers in solution. Network chains that are stretched or shrunken by the deformation of the network contribute to the elasticity of the network and thus are called elastically effective. Network chains that form loops are not stretched or shrunken by the network deformation and thus are called elastically ineffective. Loop extrusion causes the transport of chromatin segments in the elastically effective fraction of the network to the elastically ineffective loops. This means that loop extrusion stiffens the network by decreasing the number of chromatin segments in the elastically effective fraction of the network. We here take into account the stiffening of the network by the loop extrusion in an extension of the statistical thermodynamic theory of polymer gels to understand the mechanism of the assembly of sparklers. Our theory predicts that entangled DNA networks show a discontinuous transition between swollen and collapsed states − a volume phase transition,^19,20^ by changing the loop extrusion activity of condensin.

## Model

### Entangled DNA gel

We here construct a minimal model of entangled chromosomes. We treat these chromosomes as very long flexible chains that are entangled on the experimental time scale. The chains are assumed to be electrically neutral because the electric fields generated from the electric charges of DNA are screened by salt ions in the solution at physiological salt concentrations.^21^ The chromosomes are treated as an entangled DNA gel composed of *N* (Kuhn) segments of length *b*. In our model we take into account the entanglements, the loop extrusion process, and the transient crosslinks by linker histone H1.8 in an extension of the theory of polymer gels.^22^ We use the affine tube model to treat the entanglements.^15^

The free energy density (per unit volume of the final state) of the entangled DNA gel has the form

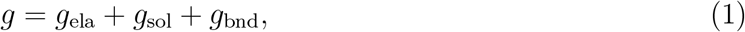

where *g*_ela_ is the elastic free energy of the network, *g*_sol_ is the solution free energy that includes the transient crosslinking by linker histones and the mixing free energy between DNA and solvent, and *g*_bnd_ is the free energy involved in the binding of linker histones to DNA. This free energy density is a function of the DNA volume fraction *ϕ* and the occupancy *α* of DNA binding sites by linker histones. The loops are produced by the loop extrusion, which is an active process, and are not densely packed at the periphery. These loops thus do not contribute to the free energy of the system.

Without the loop extrusion by condensin, all the segments in the entangled DNA network are elastically effective. The extent of entanglements between the DNA chains is defined by the number of segments *N*_e0_ between effective crosslinks due to the entanglements. With the loop extrusion process switched on, *N*_l_ segments are extruded to form loops, which are elastically ineffective. The remaining number of segments in the elastically effective part of the network is *N* − *N*_l_. With these loops, the number of segments *N*_e_ between effective crosslinks has the form

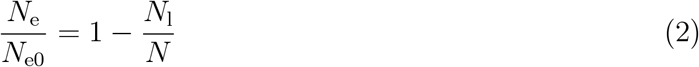

because the number *N/N*_e0_ of effective crosslinks due to the entanglements is constant. In deriving eq. (2), we used the fact that the effective crosslinks due to the entanglements can slide along the chain to share the load. The elastic energy density thus has the form

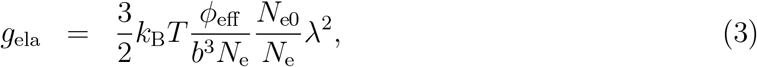

where the factor *N*_e0_*/N*_e_ represents the stiffening effect caused the loop extrusion process. *ϕ*_eff_ (= *N*_e_*ϕ/N*_e0_) is the volume fraction of elastically effective chains (the factor *ϕ*_eff_ /(*b*^3^*N*_e_) is the number density of ‘entanglement strands’ that are subchains between adjacent effective crosslinks). *k*_B_ is the Boltzmann constant and *T* is the absolute temperature. λ is the extension ratio with respect to ‘the reference state’ (discussed below). The extension ratios are equal for all the directions because we only treat the swelling and deswelling of the network and the network is isotropic. The extension ratio λ has a relationship

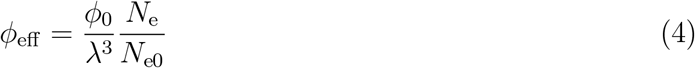

with the volume fraction *ϕ*_eff_ of elastically effective chains after the deformation, where *ϕ*_0_ is the DNA volume fraction at the reference state, see also eq. (2).

The reference state is a synthetic polymer gel where the entanglements are trapped by crosslinking. Instead of treating the entanglement process of mouse sperm chromosomes, which can be complex, we define the reference state as a DNA solution that forms the same number of effective crosslinks by entanglements as the number of entanglements in the mouse sperm chromosomes. Eq. (3) is rewritten in the form

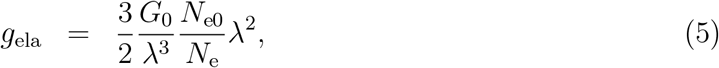

by introducing the shear modulus *G*_0_ (= *k*_B_*Tϕ*_0_/(*b*^3^*N*_e0_)) of the network at the reference state. Fortunately, although the reference state is a gedanken concept, the final result depends on the reference state only via a single parameter 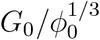, which, in principle, is experimentally accessible by measuring the force-extension relationship of entangled mouse sperm chromosomes.

The solution free energy has the form

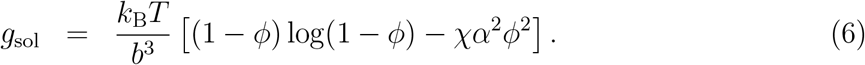

The volume fraction *ϕ* includes both DNA segments in elastically effective chains and the loops in the core, but does not include DNA segments in the loops at the surface. The first term of eq. (6) is the free energy contribution of the mixing entropy of DNA and solvent molecules. The second term of eq. (6) represents the effective interaction due to the transient crosslinking of DNA by linker histones and *χ* is the corresponding interaction parameter.^23^ This effective interaction represents the tendency for mixtures of DNA and linker histones to phase separate.^24,25^ Linker histones can be dissociated from DNA binding sites due to the translocation of the DNA by condensin. With eq. (6), we treat cases in which the time scale of linker histone binding is shorter than the time scale of the loop extrusion.

The binding free energy has the form

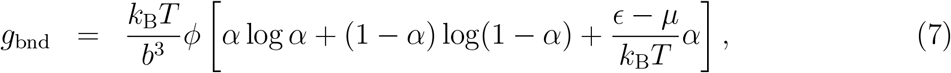

where *ϵ* is the energy due to the binding of linker histones and DNA and *μ* is the chemical potential of linker histones. Indeed, the interaction parameter has a relationship with the binding energy, 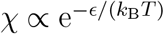.^23^

The time evolution of the DNA volume fraction *ϕ* is governed by the flux of solvent flowing in and out from the entangled DNA gels^22,26–28^ and the dynamics of binding between DNA and linker histones determine the occupancy *α*. In the steady state, the time evolution equations are reduced to the relationships

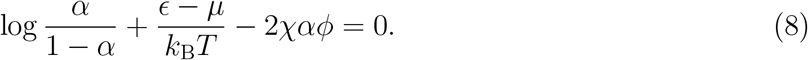

and

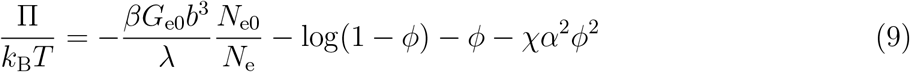

because there are no fluxes of solvent, see also refs. ^22,26–28^ for the condition of the steady state for synthetic polymer gels. One has Π = 0 for cases in which the osmotic pressure at the exterior of the entangled DNA gel is negligible. Eq. (8) implies that linker histones bind to DNA binding sites due to the binding energy *ϵ* − *μ* (the second term) and/or the interaction energy (the third term). The expression of the osmotic pressure, the right side of eq. (9), is similar to the osmotic pressure of polymer gels,^22,28^ except for the stiffening due to the loop extrusion process, see the factor *N*_e0_*/N*_e_ in the first term, and the fact that the attractive interactions between DNA segments are due to linker histones, see the factor *α*^2^ in the fourth term. The DNA volume fraction *ϕ* and the occupancy *α* are derived by using eqs. (8) and (9) once we know the fraction *N*_e0_*/N*_e_ of elastically ineffective chains, which is determined by the loop extrusion dynamics.

### Loop extrusion dynamics

We use an extension of the theory by Goloborodko and coworkers ^5^ to treat the loop extrusion process. The loop extrusion by condensin is subdivided into the loading process, the translocation process, and the unloading process. For simplicity, we here treat cases in which the loading rate of condensin molecules is relatively small and they do not form nested loops, see also Discussion. Condensin is not loaded to the DNA binding sites that are already bound by linker histones, ^29^ but can be loaded to unoccupied DNA binding sites with equal probability. In this paper, we treat cases in which the concentration of linker histones is relatively small, *ϵ* − *μ* > 0, and the binding between DNA and linker histones is stabilized by the interaction energy (the third term of eq. (8)). In such cases, the subchains at the core of the gel can be occupied by linker histones as a result of the interaction energy, while the subchains at the surface do not have enough interacting partners to be stably occupied by linker histones. The time evolution equations of the number of condensin at the core *n*_c_ and at the surface *n*_s_ has the forms

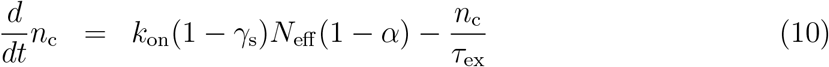

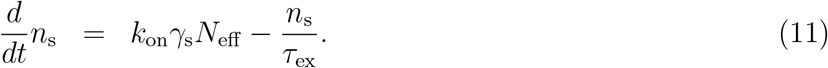

*k*_on_ is the loading rate of condensin and is proportional to the concentration of condensin in the solution. *τ*_ex_ is the average residence time of condensin. *γ*_s_ is the ratio of DNA segments at the surface. In the steady state, the numbers of DNA segments that are already translocated to the loops by condensin at the core *N*_c_ and at the surface *N*_s_ has the forms

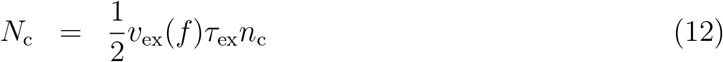

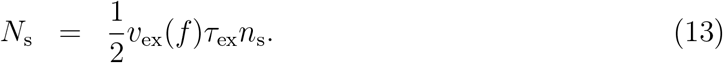

*v*_ex_(*f*) is the number of DNA segments translocated per unit time (the extrusion rate). In general, the extrusion rate *v*_ex_(*f*) depends on the tension *f* applied to the DNA. We here use the linear force-velocity relationship

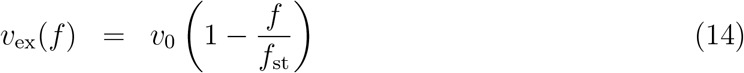

for *f* < *f*_st_ and *v*_ex_ = 0 for *f* > *f*_st_, where *v*_0_ is the extrusion rate at *f* = 0 and *f*_st_ is the stall force, analogous to the treatments for other molecular motors.^30^ The extrusion rate *v*_0_ and the stall force *f*_st_ of yeast condensin are in the order of 1.0 kpbs (∼ 3 DNA segments per second) and 1 pN, respectively, see the force-velocity relationship in ref. ^8^ The processivity *v*_0_*τ*_ex_ of condensin I is in the order of 24 kbps (∼ 80 DNA segments).^31^ With our model of entangled DNA gels, the tension *f* has the form

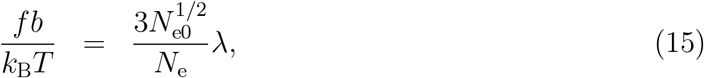

see the discussion above eq. (3).

The fraction *N*_e_*/N*_e0_ of elastically effective chains is derived by using the relationship *N*_l_ = *N*_c_ + *N*_s_ and eq. (2). The factor 1/2 in eqs. (12) and (13) represents the fact that the fraction of condensin that has translocated *l* DNA segments (0 < *l* < *vτ*_ex_) is equal at any moment of time. By solving eqs. (10) and (11) with respect to *n*_c_ and *n*_s_ in the steady state, eqs. (12) and (13) are rewritten in the forms

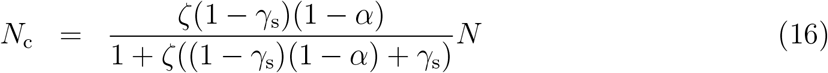

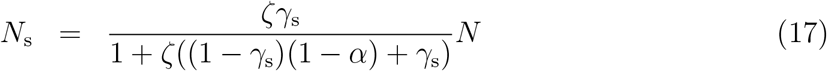

with

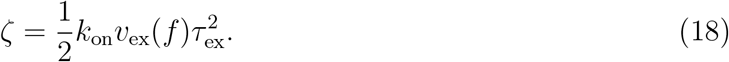

Because *ζ* is a function of the tension *f*, we define a parameter

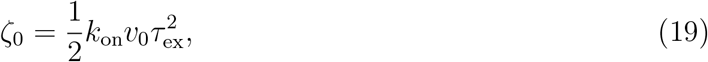

which represents the extent of the loop extrusion activity for *f* = 0.

## Results

The DNA volume fraction *ϕ* and the occupancy *α* of linker histones of the entangled DNA gel are derived by using eqs. (8) and (9) as functions of 6 parameters − the loop extrusion activity *ζ*_0_, the chemical potential of linker histones (*μ*−*ϵ*)/(*k*_B_*T*), the interaction parameter *χ*, the stall force *f*_st_*b/*(*k*_B_*T*) of condensin, the shear modulus 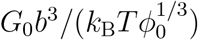 at the reference state, and the fraction *γ*_s_ of strands at the surface. One can change the loop extrusion activity *ζ*_0_ and the chemical potential *μ* of linker histones by changing the concentrations of condensin and linker histones. Eq. (8) has two stable solutions and one unstable solution of the occupancy *α* for *μ*_th1_ < *μ* < *ϵ* − 2*k*_B_*T*. The threshold chemical potential *μ*_th1_ depends only on the interaction parameter *χ* and has the analytical form

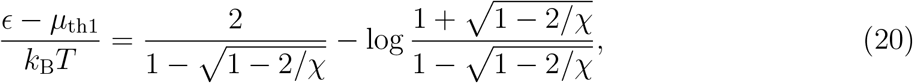

see the cyan dot in fig. 3. The derivation of eq. (20) is given in sec. S1 in the Supporting Information. The occupancy *α* is derived as the stable solution with the smaller free energy *g/ϕ*. In general, the tension developed in the chromosome can decelerate the loop extrusion of condensin. However, the essential feature of the system is captured even for the simple case that the stall force of condensin is very large, *f*_st_ → ∞, and the deceleration of the loop extrusion is negligible, *ζ* = *ζ*_0_. We thus treat this case in this section.

**Figure 2:**
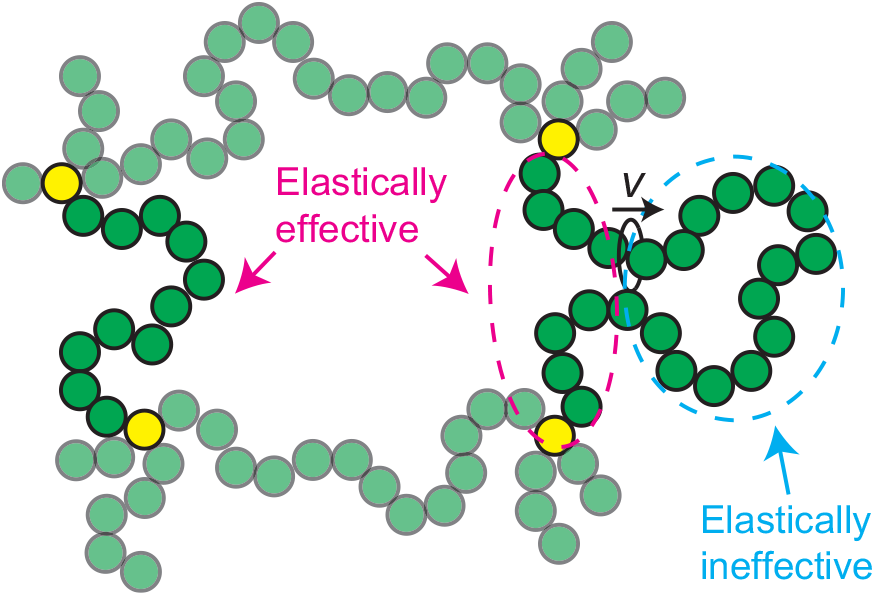
Loop extrusion of DNA in an entangled network. In the tube model, the entanglements are represented by effective crosslinks (yellow circles). The positions of the effective crosslinks displace with the deformation of the network and the chains between these crosslinks develop tension. These chains contribute to the elastic stress in the network and thus are called elastically effective, see the magenta broken line. Loops in the network are not affected by the network deformation. These loops do not contribute to the elastic stress and thus are called elastically ineffective, see the cyan broken line. The loop extrusion by condensin (black circle) translocates DNA segments (green circles) in elastically effective chains to elastically ineffective loops.

**Figure 3:**
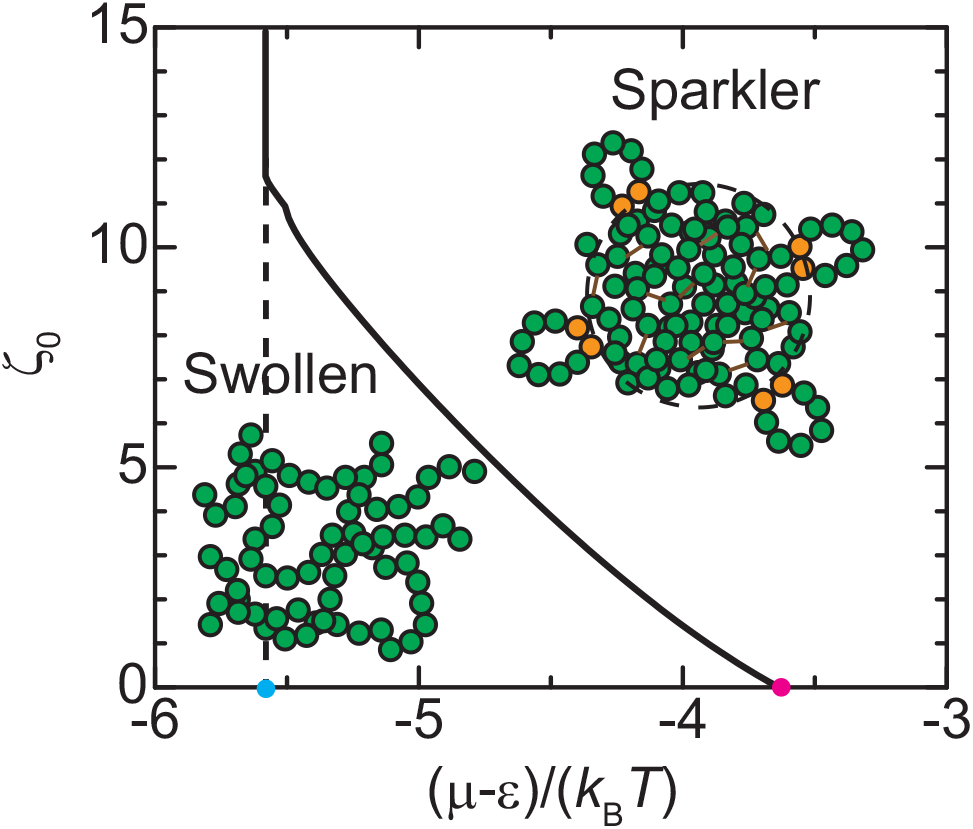
The phase diagram of an entangled DNA gel with respect to the loop extrusion activity *ζ*_0_ and the chemical potential *μ* of linker histones for *f*_st_ → ∞. The cyan and magenta dots are the two threshold values, *μ*_th1_ and *μ*_th2_, of the chemical potential. The loop extrusion activity *ζ*_0_ is defined by eq. (19). The numerical calculations are performed by using 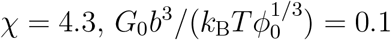, and *γ*_s_ = 0.1.

Without the loop extrusion by condensin, *ζ*_0_ = 0, the entangled DNA gel is swollen for the concentrations of linker histones in the solution smaller than a threshold value, *μ* < *μ*_th2_, whereas it collapses for *μ* > *μ*_th2_, see the magenta dot in fig. 3. The chemical potential *μ*_th2_ has the asymptotic form

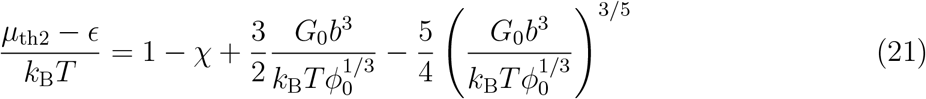

for small shear modulus, 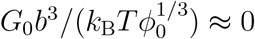, large interaction parameter, *χ* ≫ 1, and small chemical potential, (*μ* − *ϵ*)/(*k*_B_*T*) ≪ −1. Eq. (21) is derived by assuming *α* ≪ 1 and *ϕ* ≪ 1 for the swollen phase and *α* ≈ 1 and *ϕ* ≈ 1 for the collapsed state, see sec. S1 in the Supporting Information.

The DNA volume fraction *ϕ* increases with increasing the loop extrusion activity *ζ*_0_, while the occupancy *α* is very small and approximately constant for the loop extrusion activity smaller than a threshold, *ζ*_0_ *< ζ*_0th_, see the magenta and cyan lines in fig. 4. For *ζ*_0_ *< ζ*_0th_, the number of DNA segments *N*_c_ extruded to the loops in the core also increases with increasing the loop extrusion activity *ζ*_0_, see the black line in fig. 4. In the swollen phase, the DNA volume fraction *ϕ* and the number of DNA segments *N*_c_ in the core have asymptotic forms

**Figure 4:**
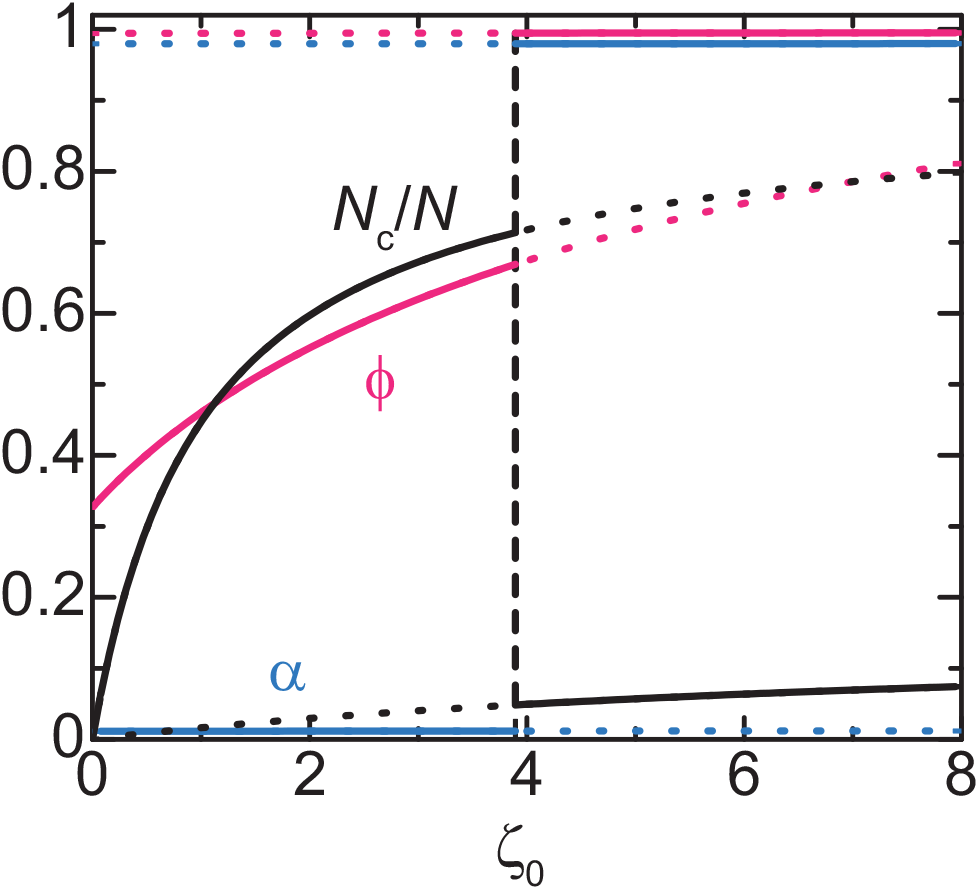
The volume phase transition of the entangled DNA gel. The volume phase transition *ϕ* (magenta), the occupancy *α* of DNA binding sites by linker histones (cyan), the fraction *N*_c_*/N* of chain segments in the loops at the core (black) are shown as a function of the loop extrusion activity *ζ*_0_ (defined by eq. (19)). These curves are derived by using *χ* = 4.3, (*μ* − *ϵ*)/(*k*_B_*T*) = −4.5, and 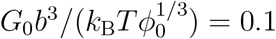, *γ*_s_ = 0.1.

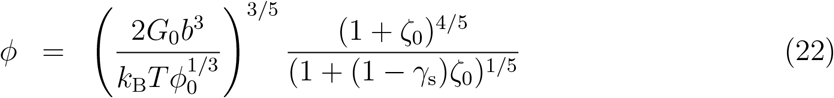

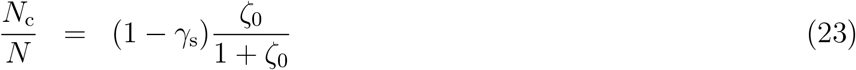

for small values of 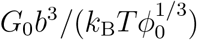, see sec. S2.1 in the Supporting Information. At the threshold loop extrusion activity, *ζ*_0_ = *ζ*_0th_, both the occupancy *α* and the DNA volume fraction *ϕ* jump to approximately unity, see the cyan and magenta lines in fig. 4, while the number *N*_c_ of DNA segments extruded to the loops in the core jumps to a smaller value, see the black line in fig. 4. The jump of the DNA volume fraction *ϕ* is analogous to the volume phase transition of polymer gels.^19,20,28^ The volume phase transition is driven by the loop extrusion process of condensin: the swollen state is destabilized by the stiffening of the entangled DNA network due to the loop extrusion process and the collapse of the network results from the transient crosslinking by linker histones. This mechanism is very different from the volume phase transition of polymer gels at thermodynamic equilibrium, where the transition is driven by the osmotic pressure of counterions^32^ or the cooperative hydration. ^33^ The small number *N*_c_ of DNA segments extruded to the loop in the core results from the fact that the loading of condensin to the DNA segments in the core is suppressed by the linker histones bound to these segments, see the black line in fig. 4 and eq. (16). Condensin is localized at the surface of the entangled DNA network. The features of the entangled DNA network in the collapsed phase are analogous to sparklers. Our theory thus predicts that sparklers are assembled by the volume phase transition of entangled chromosomes.

Now we take into account the deceleration of loop extrusion by the tension developed in the DNA. The volume phase transition of the entangled DNA gel is driven by changing the loop extrusion activity *ζ*_0_ for a window 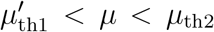 of the chemical potential of linker histones, as in the case of *f*_st_ → ∞, see fig. 5. There are two features, which are different from the case of *f*_st_ → ∞: First, the width 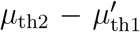 of the window of the chemical potential decreases with decreasing the stall force, see fig. 6. Second, the entangled DNA gel experiences a reentrant transition from the collapsed phase to the swollen phase by increasing the loop extrusion activity *ζ*_0_ to higher values, see fig. 5. These features imply that the sparkler phase is destabilized by the deceleration of the loop extrusion process. The reentrant volume phase transition results from the fact that the condensin is stalled at a smaller loop extrusion activity *ζ*_0_ in the swollen phase than in the collapsed phase, see the factor λ in eq. (15): the free energy of the collapsed phase increases with increasing the loop extrusion activity, while the free energy of the swollen phase, in which condensin stalls, is constant.

**Figure 5:**
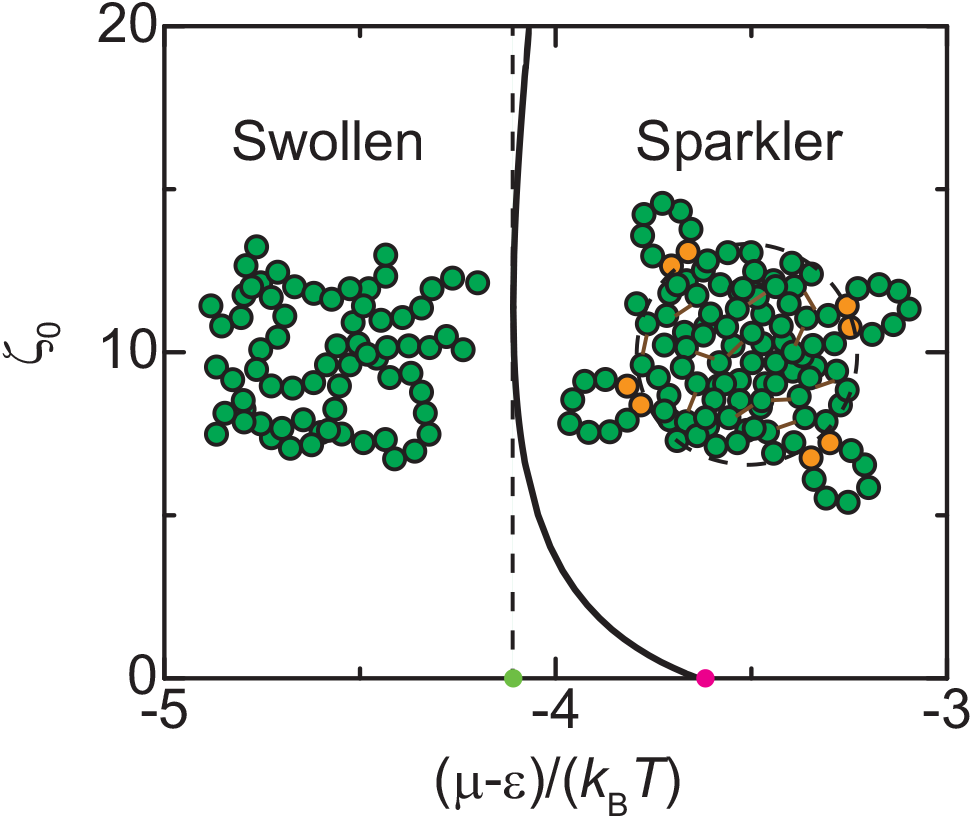
The phase diagram of an entangled DNA gel with respect to the loop extrusion activity *ζ*_0_ and the chemical potential *μ* of linker histones for *f*_st_*b/*(*k*_B_*T*) = 5.0. The light green and magenta dots are the two threshold values, 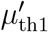 and *μ*_th2_, of the chemical potential. The loop extrusion activity *ζ*_0_ is defined by eq. (19). The numerical calculations are performed by using *χ* = 4.3, 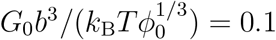, and *γ*_s_ = 0.1.

**Figure 6:**
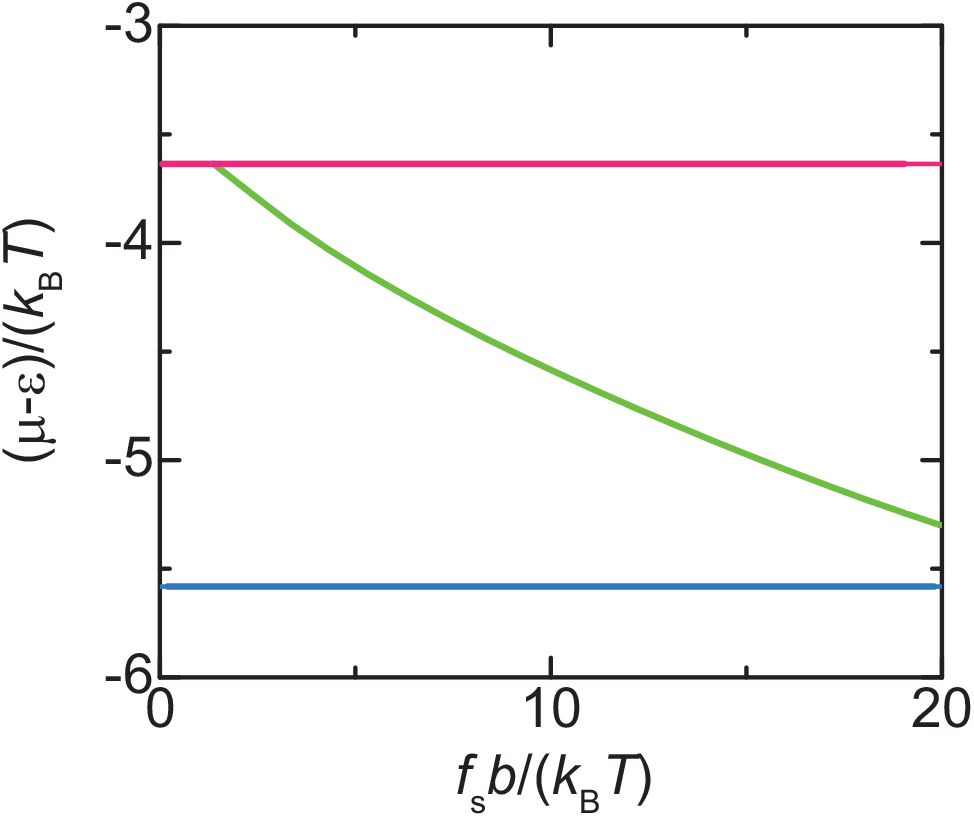
The threshold chemical potentials *μ*_th1_ (cyan), 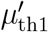 (light green), and *μ*_th2_ (magenta) vs the stall force *f*_st_. The numerical calculations are performed by using *χ* = 4.3, 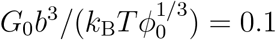, and *γ*_s_ = 0.1.

## Discussion

We have constructed a minimal model for reconstituted chromosomes, from which topo II is depleted. The reconstituted chromosomes behave as swollen entangled DNA gels if condensin is depleted. The main concept delineated by this theory is that the loop extrusion by condensin translocates DNA segments from the elastically effective chains of the entangled DNA network to the elastically ineffective chains and thus stiffens the entangled DNA network. The stiffening by the loop extrusion process destabilizes the swollen phase and drives the volume phase transition to the collapsed phase, where the attractive interaction between the DNA segments results from the transient crosslinking by linker histone H1.8. This mechanism of the volume phase transition is very different from the classical mechanisms, such as the osmotic pressure of counterions^32^ and cooperative hydration. ^33^ The linker histones that bind to the DNA segments at the core of the gel suppress the loading of condensin to these segments. Condensin is thus localized at the surface of the gel. The predicted features of the collapsed phase are analogous to the sparklers found by Shintomi and Hirano. ^14^ The sparklers typically feature a couple of protrusions. These protrusions may result from the adhesion of condensin at the surface to the cover slip and are beyond the scope of our theory. Our model also takes into account the deceleration of the loop extrusion by the tension developed in the DNA. The deceleration mechanism can be significant in entangled DNA gels, where the tension developed in DNA does not relax due to the entanglements. This situation is in contrast to the loop extrusion of DNA freely diffusing in a solution, where the tension generated by the loop extrusion relaxes relatively fast. ^34^ The stall force of yeast condensin is reported to be on the order of 1 pN, corresponding to *f*_st_*b/*(*k*_B_*T*) ≈ 24.^8^ Whether the deceleration mechanism is significant or not depends on the shear modulus 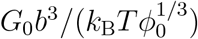 of the entangled DNA network.

We used a couple of assumptions to simplify the model. First, we assumed that condensing is not loaded to DNA loops that have been translocated by previously loaded condensin. This is the case when the concentration of condensin in the solution is relatively small. However, this assumption is not essential for our treatment: in the other limit of large condensin concentrations, the DNA segments that were translocated to the loops return to the elastically effective chains when condensin is unloaded, but are soon translocated to the loops by other condensin. Such cases can be treated by using the same formalism, but without the factor 1/2 in eqs. (12) and (13). Second, we assumed a reference state, a DNA solution that forms the same number of effective crosslinks by entanglements as the mouse sperm chromosomes. Such a reference state surely exists for cases in which chromosomes are relaxed to the thermodynamic equilibrium before topo II is depleted. However, it is not a trivial whether the reference state exists in general. Our assumption is probably the best given our current knowledge of the polymer entanglement and its simplicity.

The main prediction of our theory is the volume phase transition of entangled chromosomes by changing the loop extrusion activity for a window of the chemical potential of linker histones, see the phase diagram in fig. 5. This assumption may be experimentally accessible on reconstituted chromosomes, from which topo II is depleted, by changing the concentration of condensin and linker histones. The volume phase transition of gels of synthetic polymers is a very slow process. However, the volume phase transition of chromosomes can be fast because the chromosomes, which are on the order of micrometer, are smaller than typical synthetic polymer gels and the mesh size is likely to be rather large. It is therefore tempting to think that the volume phase transition driven by the loop extrusion process is involved in the condensation of chromosomes at the entry of mitosis. Indeed, Beel and coworkers recently proposed a similar view, but with the volume phase transition due to the osmotic pressure of counterions.^32^ Whether the volume phase transition or the disentanglement by topo II are faster remains to be understood and will be subject of our future research. At least, our theory provides insight into the mechanism of the assembly of the reconstituted chromosomes, which have been studied intensively in recent years. ^12–14,29,32^

## Acknowledgement

This work was supported by JSPS KAKENHI Grant Number 20H05934 (Genome Modality), 21K03479, 21H00241. T.Y. acknowledges discussions with T. Hirano (Riken), K. Shintomi (Riken), H. Niki (National Institute for Genetics), and T. Sakaue (Aoyama Gakuin). H.S. was supported by the Deutsche Forschungsgemeinschaft (DFG, German Research Foundation) under Germany’s Excellence Strategy – EXC-2068 – 390729961

## Supporting Information Available

Supporting Information: Sec. S1 Derivation of threshold chemical potential *μ*_th1_ and sec. S2 Approximate solution.

