## Supporting Information for "Loop extrusion driven volume phase transition of entangled chromosomes"

### S1 Derivation of threshold chemical potential $\mu_{\rm th1}$

The equality of the chemical potential of linker histones is given by

$$\log\frac{\alpha}{1-\alpha}+\frac{\epsilon-\mu}{k_{\rm B}T}-2\chi\alpha\phi=0. \tag{S1}$$

see also eq. (8) in the main article. Eq. (S1) is rewritten in the form

$$2\chi\phi=\frac{1}{\alpha}\left(\log\frac{\alpha}{1-\alpha}+\frac{\epsilon-\mu}{k_{\rm B}T}\right). \tag{S2}$$

The first and second derivatives of eq. (S2) have the forms

$$\frac{\partial}{\partial\alpha}(2\chi\phi) = -\frac{1}{\alpha^2}\left(\log\frac{\alpha}{1-\alpha}+\frac{\epsilon-\mu}{k_{\rm B}T}-\frac{1}{1-\alpha}\right) \tag{S3}$$

$$\frac{\partial^2}{\partial\alpha^2}(2\chi\phi) = \frac{2}{\alpha^3}\left(\log\frac{\alpha}{1-\alpha}+\frac{\epsilon-\mu}{k_{\rm B}T}-\frac{1}{1-\alpha}-\frac{1}{2}\frac{1-2\alpha}{(1-\alpha)^2}\right). \tag{S4}$$

For simplicity, we first release the condition $\phi\leq 1$. The condition with which eq. (S2) has two stable solutions and one unstable solution is that $\frac{\partial}{\partial\alpha}(2\chi\phi)\leq 0$ at the value of the occupancy $\alpha$, such that $\frac{\partial^2}{\partial\alpha^2}(2\chi\phi)=0$. The critical condition is $\frac{\partial}{\partial\alpha}(2\chi\phi)=\frac{\partial^2}{\partial\alpha^2}(2\chi\phi)=0$. This leads to $\alpha=1/2$ and $(\epsilon-\mu)/(k_{\rm B}T)=2$. The condition with which eq. (S2) has multiple solutions is therefore

$$\frac{\epsilon-\mu}{k_{\rm B}T}\geq 2. \tag{S5}$$

For $(\epsilon-\mu)/(k_{\rm B}T)>2$, there are two solutions of $\frac{\partial}{\partial\alpha}(2\chi\phi)=0$, $\alpha_{\rm max}$ and $\alpha_{\rm min}$ ($\alpha_{\rm max}<\alpha_{\rm min}$). The solution $\alpha_{\rm max}$ corresponds to the local maximum of $\phi$, whereas the solution $\alpha_{\rm min}$ corresponds to the local minimum of $\phi$. When the local minimum of $\phi$ is larger than 1, the solution is unique. The condition with which the local minimum of $\phi$ is 1 is that $\phi=1$ and $\frac{\partial}{\partial\alpha}(2\chi\phi)=0$. The occupancy $\alpha$ and the chemical potential $(\epsilon-\mu_{\rm th1})/(k_{\rm B}T)$ with this condition are

$$\alpha_{\rm min} = \frac{1}{2} + \sqrt{\frac{1}{4} - \frac{1}{2\chi}} \tag{S6}$$

$$\frac{\epsilon - \mu_{\rm th1}}{k_{\rm B}T} = \frac{2}{1 - \sqrt{1 - 2/\chi}} - \log \frac{1 + \sqrt{1 - 2/\chi}}{1 - \sqrt{1 - 2/\chi}}. \tag{S7}$$

There are therefore two stable solutions and one unstable solution for $\mu_{\rm th1} < \mu < \epsilon - 2k_{\rm B}T$.

### S2 Approximate solution

The free energy density and the osmotic pressure of the system are given in eqs. (1) and (9) in the main article and have the form

$$\begin{aligned} \frac{gb^3}{\phi k_{\rm B}T} &= \frac{3}{2}\frac{G_{\rm e0}b^3}{\phi_0^{1/3}k_{\rm B}T}\frac{N_{\rm e0}/N_{\rm e}}{(1 - N_{\rm s}/N)^{1/3}}\phi^{-2/3} + \frac{1-\phi}{\phi}\log(1-\phi) - \chi\alpha^2\phi \\ &\quad +\alpha\log\alpha + (1-\alpha)\log(1-\alpha) + \frac{\epsilon - \mu}{k_{\rm B}T}\alpha + \frac{\Pi b^3}{\phi k_{\rm B}T} \end{aligned} \tag{S8}$$

$$\frac{\Pi b^3}{k_{\rm B}T} = -\frac{G_{\rm e0}b^3}{\phi_0^{1/3}k_{\rm B}T}\frac{N_{\rm e0}/N_{\rm e}}{(1 - N_{\rm s}/N)^{1/3}}\phi^{1/3} - \log(1-\phi) - \phi - \chi\alpha^2\phi^2. \tag{S9}$$

The numbers of DNA segments, $N_{\rm c}$ and $N_{\rm s}$, in the loops at the interior and the surface have the forms

$$N_{\rm c} = \frac{\zeta(1-\gamma_{\rm s})(1-\alpha)}{1 + \zeta((1-\gamma_{\rm s})(1-\alpha) + \gamma_{\rm s})}N \tag{S10}$$

$$N_{\rm s} = \frac{\zeta\gamma_{\rm s}}{1 + \zeta((1-\gamma_{\rm s})(1-\alpha) + \gamma_{\rm s})}N \tag{S11}$$

in the steady state with

$$\zeta = \zeta_0\left(1 - \frac{f}{f_{\rm st}}\right). \tag{S12}$$

The fraction of DNA segments in elastically effective chains has the form

$$\frac{N_{\rm e}}{N_{\rm e0}} = 1 - \frac{N_{\rm c} + N_{\rm s}}{N}, \tag{S13}$$

see eqs. (16) - (19) in the main article. $\gamma_{\rm s}$ is the fraction of DNA segments at the surface. $\zeta_0$ is the loop extrusion activity defined in eq. (19) in the main article. $f_{\rm st}$ is the stall force of condensin. $f$ is the tension applied to the elastically effective DNA chains and has the form

$$\frac{fb}{k_{\rm B}T} = \frac{3N_{\rm e0}^{1/2}}{N_{\rm e}}\lambda. \tag{S14}$$

#### S2.1 Swollen phase

The occupancy $\alpha$ of linker histones and the DNA volume fraction $\phi$ are small in the swollen phase unless the loop extrusion activity is very large, see fig. 4 in the main article. We here derive an approximate solution $\alpha = \alpha_1$ and $\phi = \phi_1$ ($\alpha_1 \ll 1$ and $\phi_1 \ll 1$). By first order with respect to $\alpha_1$ and $\phi_1$, eqs. (S1), (S8), and (S9) is approximated in the forms

$$\log \alpha_1 + \frac{\epsilon - \mu}{k_{\rm B}T} \simeq 0 \tag{S15}$$

$$\frac{\Pi b^3}{k_{\rm B}T} \simeq -\frac{G_{\rm e0}b^3}{\phi_0^{1/3}k_{\rm B}T}\frac{(1+\zeta)^{4/3}}{(1+(1-\gamma_{\rm s})\zeta)^{1/3}}\phi_1^{1/3} + \frac{1}{2}\phi_1^2 \tag{S16}$$

$$\frac{gb^3}{\phi_1 k_{\rm B}T} \simeq -1 + \frac{5}{4}\phi_1. \tag{S17}$$

$\alpha_1$ and $\phi_1$ are derived in the forms

$$\alpha_1 = {\rm e}^{-(\epsilon-\mu)/(k_{\rm B}T)} \tag{S18}$$

$$\phi_1 = \left(\frac{2G_{\rm e0}b^3}{\phi_0^{1/3}k_{\rm B}T}\right)^{3/5}\frac{(1+\zeta)^{4/5}}{(1+(1-\gamma_{\rm s})\zeta)^{1/5}} \tag{S19}$$

by using eqs. (S15) and (S16) for $\Pi = 0$. Eq. (S19) is equal to eq. (22) in the main article. The approximate forms of the fractions of DNA segments in elastically effective chains and in the loops at the surface are derived as

$$\frac{N_{\rm e}}{N_{\rm e0}} \simeq \frac{1}{1+\zeta} \tag{S20}$$

$$\frac{N_{\rm s}}{N} \simeq \frac{\zeta}{1+\zeta}\gamma_{\rm s} \tag{S21}$$

by using eqs. (S10) and (S11).

#### S2.2 Sparkler phase

The occupancy $\alpha$ of linker histones and the DNA volume fraction $\phi$ are approximately unity in the sparkler phase. We here derive the approximate solutions by assuming $\alpha = 1 - \delta\alpha_2$ and $\phi = 1 - \delta\phi_2$ ($\delta\alpha_2 \ll 1$ and $\delta\phi_2 \ll 1$). By the first order with respect to $\delta\alpha_2$ and $\delta\phi_2$, eqs. (S1), (S8), and (S9) are approximated by the forms

$$-\log\delta\alpha_2 + \frac{\epsilon-\mu}{k_{\rm B}T} - 2\chi = 0. \tag{S22}$$

$$\frac{\Pi b^3}{k_{\rm B}T} \simeq -\frac{G_{\rm e0}b^3}{\phi_0^{1/3}k_{\rm B}T}(1+\gamma_{\rm s}\zeta)^{4/3} - \log\delta\phi_2 - 1 - \chi \tag{S23}$$

$$\frac{gb^3}{k_{\rm B}T} \simeq \frac{3}{2}\frac{G_{\rm e0}b^3}{\phi_0^{1/3}k_{\rm B}T}(1+\gamma_{\rm s}\zeta)^{4/3} - \chi + \frac{\epsilon-\mu}{k_{\rm B}T}. \tag{S24}$$

$\delta\alpha_2$ and $\delta\phi_2$ are derived in the forms

$$\delta\alpha_2 = {\rm e}^{(\epsilon-\mu)/(k_{\rm B}T)-2\chi}. \tag{S25}$$

$$\delta\phi_2 = {\rm e}^{-1-\chi-\frac{G_{\rm e0}b^3}{\phi_0^{1/3}k_{\rm B}T}(1+\gamma_{\rm s}\zeta)^{4/3}}. \tag{S26}$$

by using eqs. (S22) and (S23) for $\Pi = 0$. The approximate expressions of the fractions of DNA segments in elastically effective chains and in the loops at the surface are derived as

$$\frac{N_{\rm e}}{N_{\rm e0}} \simeq \frac{1}{1+\zeta\gamma_{\rm s}} \tag{S27}$$

$$\frac{N_{\rm s}}{N} \simeq \frac{\zeta\gamma_{\rm s}}{1+\zeta\gamma_{\rm s}}, \tag{S28}$$

by using eq. (S10) and (S11).

##### S2.3 Swollen-sparkler transition for $\zeta \simeq \zeta_0$

We here treat cases in which the deceleration of the loop extrusion by the tension applied to DNA is not significant, $\zeta \simeq \zeta_0$. The loop extrusion activity at the transition between swollen phase and sparkler phase is derived by the equality of free energy between the two phases, see eqs. (S17) and (S24). This leads to the chemical potential $\mu_{\rm th}$ at the transition in the form

$$\frac{\epsilon - \mu_{\rm th}}{k_{\rm B}T} = \chi - 1 + \frac{5}{4}\left(\frac{2G_{\rm e0}b^3}{\phi_0^{1/3}k_{\rm B}T}\right)^{3/5} \frac{(1+\zeta_0)^{4/5}}{(1+(1-\gamma_{\rm s})\zeta_0)^{1/5}} - \frac{3}{2}\frac{G_{\rm e0}b^3}{\phi_0^{1/3}k_{\rm B}T}(1+\gamma_{\rm s}\zeta_0)^{4/3}. \tag{S29}$$

The threshold chemical potential $\mu_{\rm th2}$ at which the transition between the two phases happens for $\zeta_0 \to 0$ and thus has the form

$$\frac{\epsilon - \mu_{\rm th2}}{k_{\rm B}T} \simeq \chi - 1 - \frac{3}{2}\frac{G_{\rm e0}b^3}{\phi_0^{1/3}k_{\rm B}T} + \frac{5}{4}\left(\frac{G_{\rm e0}b^3}{\phi_0^{1/3}k_{\rm B}T}\right)^{3/5}. \tag{S30}$$
